# A machine learning classifier trained on cancer transcriptomes detects NF1 inactivation signal in glioblastoma

**DOI:** 10.1101/075382

**Authors:** Gregory P. Way, Robert J. Allaway, Stephanie J. Bouley, Camilo E. Fadul, Yolanda Sanchez, Casey S. Greene

## Abstract

**Background:** We have identified molecules that exhibit synthetic lethality in cells with loss of the neurofibromin 1 (*NF1*) tumor suppressor gene. However, recognizing tumors that have inactivation of the *NF1* tumor suppressor function is challenging because the loss may occur via mechanisms that do not involve mutation of the genomic locus. Degradation of the NF1 protein, independent of *NF1* mutation status, photocopies inactivating mutations to drive tumors in human glioma cell lines. NF1 inactivation may alter the transcriptional landscape of a tumor and allow a machine learning classifier to detect which tumors will benefit from synthetic lethal molecules.

**Results:** We developed a strategy to predict tumors with low NF1 activity and hence tumors that may respond to treatments that target cells lacking NF1. Using RNAseq data from The Cancer Genome Atlas (TCGA), we trained an ensemble of 500 logistic regression classifiers that integrates mutation status with whole transcriptomes to predict NF1 inactivation in glioblastoma (GBM). On TCGA data, the classifier detected *NF1* mutated tumors (test set area under the receiver operating characteristic curve (AUROC) mean = 0.77, 95% quantile = 0.53 – 0.95) over 50 random initializations. On RNA-Seq data transformed into the space of gene expression microarrays, this method produced a classifier with similar performance (test set AUROC mean = 0.77, 95% quantile = 0.53 – 0.96). We applied our ensemble classifier trained on the transformed TCGA data to a microarray validation set of 12 samples with matched RNA and NF1 protein-level measurements. The classifier’s NF1 score was associated with NF1 protein concentration in these samples.

**Conclusions:** We demonstrate that TCGA can be used to train accurate predictors of NF1 inactivation in GBM. The ensemble classifier performed well for samples with very high or very low NF1 protein concentrations but had mixed performance in samples with intermediate NF1 concentrations. Nevertheless, high-performing and validated predictors have the potential to be paired with targeted therapies and personalized medicine.

## BACKGROUND

Genomic tools allow investigators to devise therapies targeting specific molecular abnormalities in tumors. One such alteration is the loss of neurofibromin 1 (NF1), an important tumor suppressor that regulates the activity of *RAS* GTPases [1,2]. Heterozygous mutation or deletion of *NF1* causes neurofibromatosis type 1 (NF), one of the most frequently inherited genetic disorders [3]. NF patients often develop plexiform neurofibromas (PNs), benign nerve tumors for which the only therapy is surgery. However, resection is often impossible due to the tumor’s intimate association with peripheral and cranial nerves [4]. PNs can transform to malignant peripheral nerve sheath tumors (MPNSTs), which are chemo-and radiation-resistant sarcomas with a dismal 20% 5-year survival [5]. In addition, patients with NF are susceptible to a broad spectrum of other tumors including low-grade/pilocytic astrocytomas, pheochromocytomas, optic nerve gliomas, and juvenile myelomonocytic leukemias [6]. Many aggressive non-NF associated (sporadic) tumors have recently been shown to harbor *NF1* mutations, including glioblastoma (GBM), neuroblastoma, melanoma, thyroid, ovarian, breast, and lung cancers [7]. Therefore, somatic and inherited loss of *NF1* function is emerging as a driver of tumors from different organ sites.

Several groups including our own have been working to develop therapeutic approaches to target tumors with loss of NF1. Previously, our lab developed a high throughput approach using yeast and mammalian screening platforms to identify tool compounds and drug targets for cancer cells in which NF1 loss drives tumor formation. Our pipeline identified small molecules that selectively kill or stop the growth of MPNST cells carrying a mutation in *NF1* or yeast lacking the *NF1* homolog *IRA2* [8]. We also developed an assay in yeast to identify the targets of our lead tool compounds and found that one of these compounds (UC-1) shares a mechanism (phosphorylation of RNA Pol II CTD Ser2/5) with experimental drugs in clinical trials [8]. UC-1 impacts CTD phosphorylation, which is regulated by the CTD kinase Ctk1, the yeast homolog of human Cdk9. We showed that deletion of *CTK1* was synthetic lethal with loss of the yeast *NF1* homolog *IRA2*. Furthermore, we have found that inhibitors of this process (dinaciclib, SNS-032) can inhibit other types of RAS-dysregulated tumor cells [9].

However, relying on genetic data alone to identify tumors that may be susceptible to therapies targeting NF1 loss may leave a proportion of potentially actionable tumors unrecognized. NF1 tumor suppressor activity can be lost via mutation of the genomic locus, proteasome-mediated degradation, inhibition by miRNA, *de novo* insertion of an ALU element, and C→U editing of the *NF1* mRNA [10–14]. This complexity presents challenges when trying to identify tumors that will benefit from molecules that exert synthetic lethality with dysregulation of *NF1/RAS* pathways.

The Cancer Genome Atlas (TCGA) has released a large volume of data on several cancer tissues measured on a variety of genomic platforms. In the present study, we leverage TCGA GBM RNAseq expression data with matched mutation calls to construct a classifier capable of identifying an NF1 inactivation signature. This strategy sidesteps the problem of functional characterization of mutations by evaluating a regulator’s downstream gene expression activity. We applied this signature to predict NF1 inactivation in a cohort of biobanked GBMs. In general, this approach can be translatable to any gene producing measurable downstream transcriptome-wide effects.

## METHODS

### The Cancer Genome Atlas Data used for building the classifier

We downloaded RNAseq and mutation data from TCGA Pan Cancer project from the UCSC Xena data portal [15] and subset each dataset to only the GBMs [16]. The data consists of 607 GBMs; of which 291 have mutation calls, 172 have RNAseq measurements, and 149 have both RNAseq and mutation calls. Of these 149 samples, 15 have inactivating *NF1* mutations (10.1%) and were used as gold standard positives in building the classifier (Supplementary Table S1). Additionally, to reduce dimensionality while avoiding unexpressed and invariant genes, we subset to the top 8,000 most variably expressed genes by median absolute deviation. We z-scored all gene expression measurements. This resulted in the final input matrix with dimension 149 samples by 8,000 genes. For use in platform independent predictions, we used Training Distribution Matching (TDM) to transform the TCGA RNAseq data to match a microarray expression distribution [17].

Since we are also aware of the *NF1* mutation status for each of the samples, we form a supervised learning task - predicting when a sample has loss of *NF1* activity. Our “X” matrix is formed by the RNAseq measurements for all 149 samples measured by 8,000 genes, which are the features in the model. Our “y” vector consist of {0, 1} elements where a 1 corresponds to a sample with an inactivating *NF1* mutation and a 0 is an *NF1* wildtype sample. The machine learning task is to find the feature weights, or gene coefficients, that best minimize our objective function. Along with these feature weights corresponding to the genes’ importance in the learning task, we also output a probability estimate for each sample that they have loss of *NF1* activity.

### Hyperparameter optimization of the logistic regression classifier

Using the GBM RNAseq data, we trained logistic regression classifiers with an elastic net penalty using stochastic gradient descent to detect tumors with NF1 inactivation. We chose a penalized regression model because it is simple to train and has easily interpretable outputs including importance scores for each gene (feature weights) associated with the downstream consequences of *NF1* loss of function and a probability for each sample that *NF1* is lost. An elastic net logistic regression model has also been successfully implemented in similar studies [18–20].

We identified high-performing alpha and L1 mixing parameters using 5-fold cross validation ensuring balanced membership of *NF1* mutations in each fold. Briefly, alpha controls how weight penalty and the L1 mixing parameter tunes the amount of test set regularization by controlling the sparsity of the features. An L1 mixing parameter value of zero corresponds to the L2 penalty and a value of one corresponds to the L1 penalty, with L1 bringing a sparser solution. We used python 3.5.1 and Sci-kit Learn for machine learning implementations [21].

### Ensemble classifier construction and application to the validation set

After selecting optimal hyperparameters, we constructed 500 classifiers that would compose our ensemble model. Specifically, across 100 different random initializations, we subset the full TCGA GBM data into 5 folds and trained a single classifier for each training fold.

We borrowed terminology from the epidemiology field to describe data partitioning. We trained our models on a “training” partition and assessed model performance on a “test” partition, which refers to the held out cross-validation fold. The independent “validation set” refers to the GBM dataset generated in a different lab (see Figure 1A).

**Figure 1.**
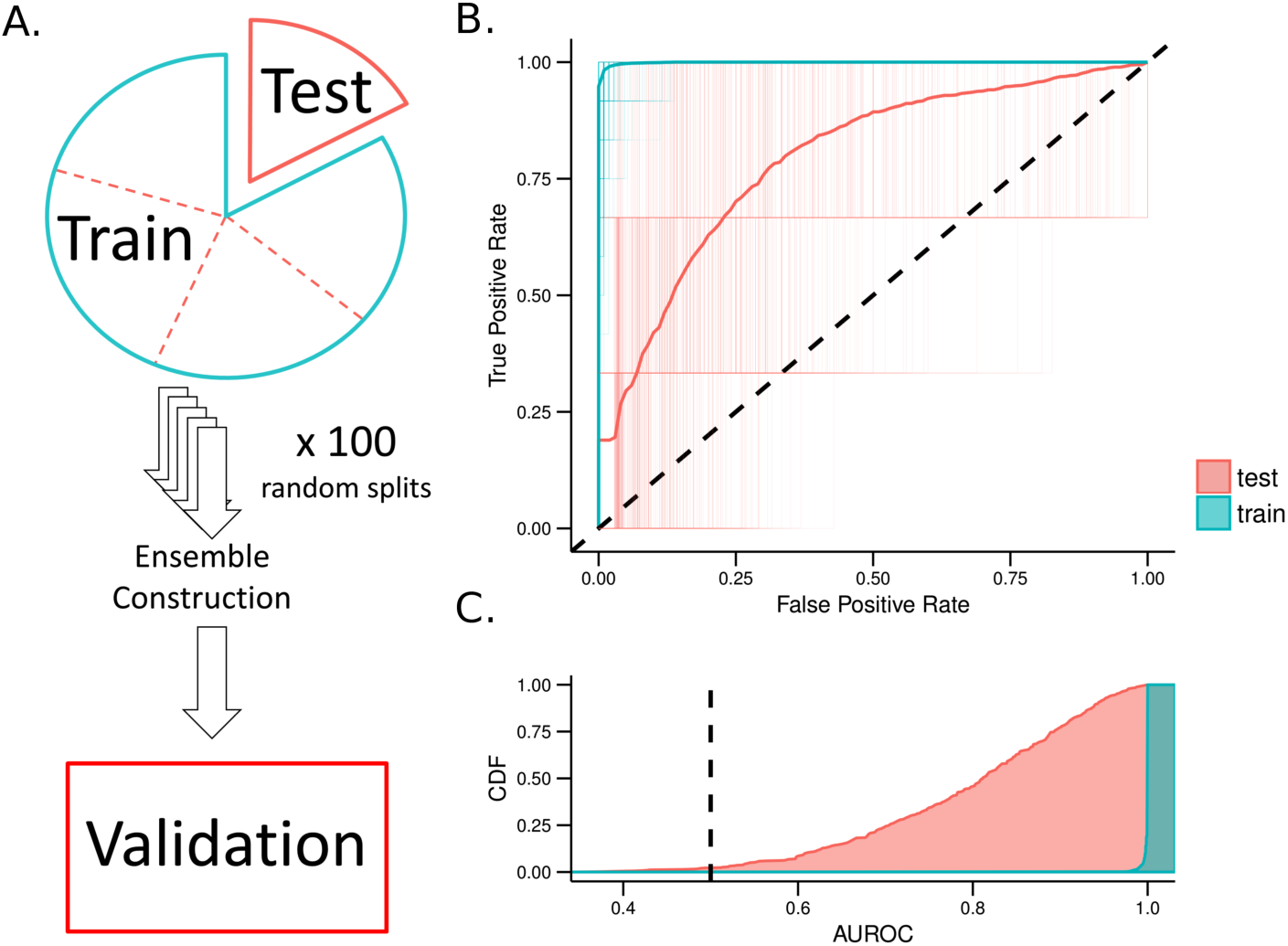
*Logistic regression classifier with elastic net penalty training and testing errors over 100 iterations for Training Distribution Matching (TDM) transformation of The Cancer Genome Atlas Glioblastoma RNAseq data*. (A) Schematic describing the terms used for training, testing, and validating our model. We applied 5-fold cross validation to the full dataset which consists of training and testing splits in each fold. The model is then applied as an ensemble classifier on a set of in-house samples (validation set) (B) Receiver operating characteristic (ROC) curves for all 500 classifiers that make up the ensemble model applied to both training and testing set. Also shown is the aggregate performance of the ensemble classifier. (C) The cumulative density of area under the ROC curve (AUROC) for training and testing partitions.

Because of the small number of gold standard positive training examples, we were concerned about the stability of our model solutions. Therefore, we constructed an ensemble classifier from the 500 models. Specifically, we assigned each classifier a weight using the specific randomization’s “test set” cross-validation AUROC. Lastly, for the final *NF1* inactivation prediction, we used the mean of the weighted predictions across all iterations as the *NF1* inactivation prediction. We applied this ensemble classifier to the validation set in which NF1 protein levels were directly measured.

### Effect sizes and power analysis

We calculated the decision function of each ensemble classifier applied to all samples in the training and testing 5-fold cross validation folds to calculate Cohen’s D effect size between predicted NF1 wildtype and NF1 inactive samples [22]. The Cohen’s D metric quantifies the difference between NF1 wildtype and NF1 inactive samples according to the mean classifier score and directly demonstrates how different the ensemble model predicts the two groups to be.

Moreover, we were also concerned that our relatively small validation set would not provide us with enough power to observe a detectable effect in the ensemble model’s final prediction. We performed a one-tailed Welch’s two-sample *t*-test comparing the NF1 protein concentration of our validation samples that were predicted to be either NF1 wildtype or NF1 deficient. Using the given sample size, Cohen’s D effect size, and a significance threshold of α = 0.05, we calculated the power of the prediction scores on the validation set. The power analysis was two-sample, one-tailed and incorporated unequal sample sizes in each group.

### Validation Sample Acquisition

Thirteen flash-frozen, de-identified GBM samples were obtained from the Maine Medical Center Biobank. Samples were received on dry ice and stored at −80°C until isolation of DNA/RNA/protein. To isolate DNA, tumor fragments of approximately 20 mg in mass were harvested on an aluminum block pre-chilled on dry ice. Samples were then immediately transferred to a mortar and pestle containing a small volume of liquid nitrogen. The fragments were pulverized in the mortar and pestle, and the liquid nitrogen was allowed to evaporate. Next, samples were immediately processed with a DNA/RNA/Protein Purification Plus kit (Norgen Biotek) following the standard operating protocol for animal tissue. DNA concentration and quality were assessed using an ND-1000 (Nanodrop), a Qubit Fluorometer (Thermo Scientific), and a Fragment Analyzer (Advanced Analytical Technologies). To isolate RNA, −80 C tumor fragments were placed in 5-10 volumes of RNAlater-ICE Frozen Tissue Transition Solution (Ambion) and placed at −20°C until RNA extraction with a mirVana miRNA isolation kit, without phenol, following the standard operating protocol (Thermo Scientific). Samples were homogenized using a manual homogenizer in the presence of mirVana lysis buffer. RNA concentration and quality were determined using a Qubit Fluorometer (Thermo Scientific) and a Fragment Analyzer (Advanced Analytical Technologies). To isolate protein, small tumor fragments were pulverized and lysed in approximately 3 volumes of ice-cold radioimmunoprecipitation assay (RIPA) buffer (150 mM sodium chloride, 1% v/v nonidet P40, 0.5% w/v sodium deoxycholate, 0.05% w/v sodium dodecyl sulfate, 50 mM Tris pH 8.0) containing 1 mM sodium orthovanadate, 1 mM sodium fluoride, 1 mM phenylmethylsulfonyl fluoride, and 1X protease inhibitor cocktail (0.1 μg/mL leupeptin, 100 μM benzamidine HCl, 1 μM aprotinin, 0.1 μg/mL soybean trypsin inhibitor, 0.1 μg/mL pepstatin, 0.1 μg/mL antipain). Samples were passed through a 25⅘ g needle and subsequently sonicated on ice to promote efficient lysis and DNA shearing. After a 30 minute incubation on ice, lysates were cleared by centrifuging at 16100 × g for 20 minutes. HEK293T, U87-MG, and U87-MG cells treated for two hours with 1 micromolar bortezomib (Selleckchem) and 10 micromolar MG132 (Selleckchem) were also prepared in RIPA buffer. Protein samples were stored at −80°C until analysis.

### Cell Culture

U87-MG and HEK293T cells were purchased from ATCC. Cell lines were regularly passaged and were cultured in Dulbecco’s Modified Eagle Medium (Corning) with 10% v/v fetal bovine serum (Gibco) at 37°C in 5% CO_2_.

Recent data regarding the U87MG cell line published by Allen *et al* suggest that the U87MG cell line distributed by ATCC is not from the same tumor as the cell line that was originally isolated in Uppsala. Transcriptome analysis comparing ATCC U87MG cell line to known tumor transcriptomes indicate that the ATCC U87MG cell line is a central nervous system tumor and is likely a glioblastoma cell line [23].

In the present study, we employ this cell line as a control representing an NF1-deficient tumor cell line. Previous studies have shown that the U87MG cell line has elevated proteasome-mediated degradation of NF1 and that this cell line required the loss of NF1 protein to promote tumorigenesis in xenograft tumor models [10]. Given that the ATCC U87MG cell line is a well-characterized and broadly-used model of NF1 deficient tumor cells [10,24–26], we propose that the use of the ATCC U87MG cell line is an appropriate control for Figure 2.

**Figure 2.**
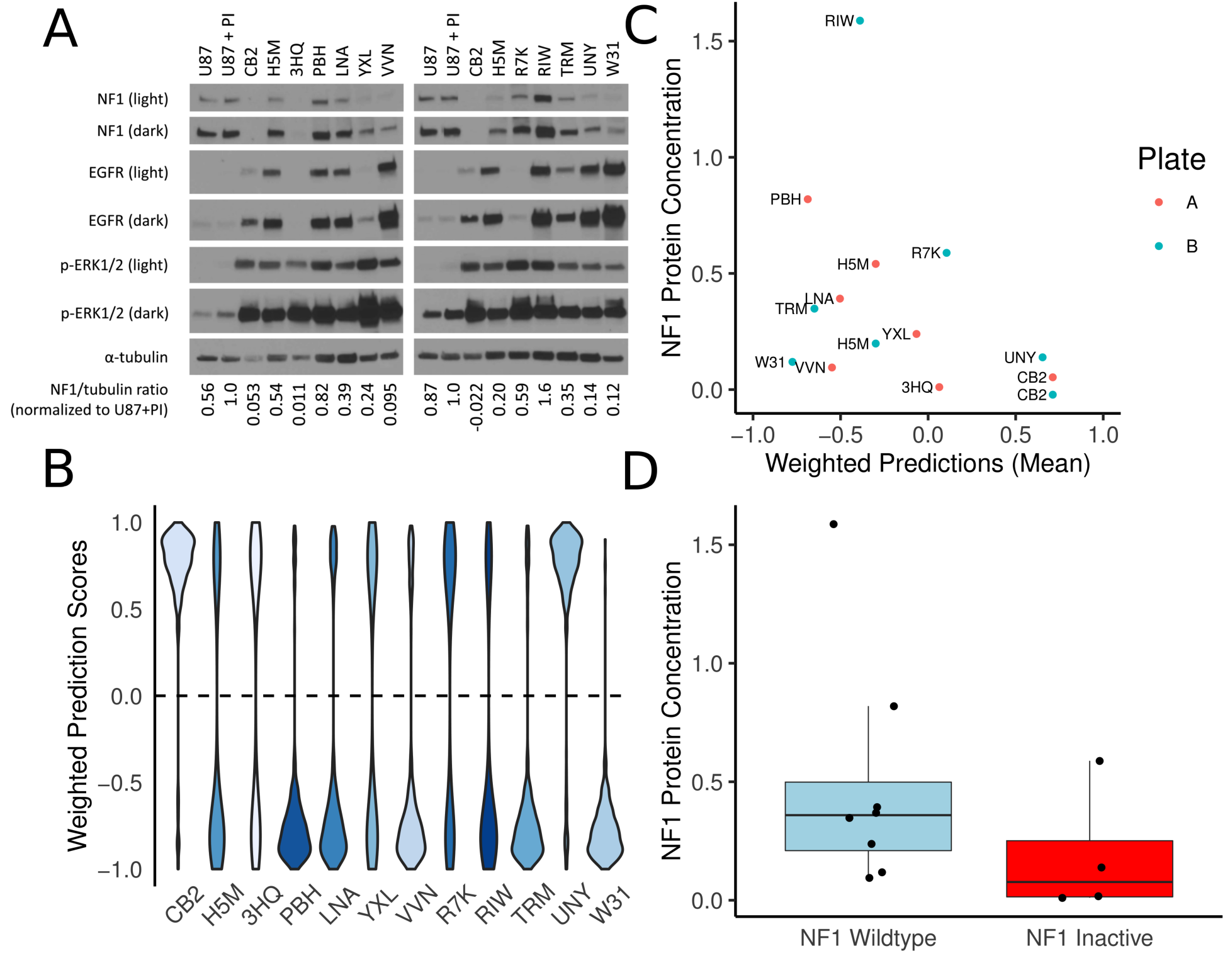
*Performance of our classifier on a validation set*. (A) Two distinct western blots for each of our twelve samples. The controls are U87-MG, an *NF1* WT glioblastoma cell line that exhibits proteasomal degradation of the NF1 protein. U87+PI are U87-MG cells are treated with the proteasome inhibitors (PI) MG-132 and bortezomib to block proteasome-mediated degradation of NF1. We used the NF1/tubulin ratio normalized to U87+PI as our NF1 protein level estimate. (B) Prediction scores for each of the 500 classifiers weighted by cross validation test set AUROC where a negative number indicates *NF1* wildtype and a positive number is indicates NF1 inactivation. Increasing grayscale indicates higher observed NF1 protein concentrations. (C) We quantify protein against U87+PI and provide the mean of the weighted predictions. (D) Based on weighted predictions, we show the abundance of NF1 protein compared to U87+PI.

### RNA Microarray

After RNA isolation and QC, samples were labeled for the GeneChip Human Transcriptome Array 2.0 (HTA 2.0, Affymetrix). Labeling was performed with Affymetrix Proprietary DNA Label (biotin-linked) using a WT Plus Kit (Affymetrix) provided with the HTA 2.0, following the standard operating protocol for HTA 2.0, including PolyA controls. Hybridization, washing, and staining were performed with the WT Plus Kit, following the standard operating protocol for HTA 2.0. Washing and staining were performed using a GeneChip Fluidics 450. Scanning was performed with a GeneChip Scanner 3000. These data were deposited in the Gene Expression Omnibus under accession GSE85033.

### Validation Sample Processing

We applied a quality control pipeline [27] to all CEL files generated by the HTA 2.0. All validation samples passed processing quality control, which included an inspection of spatial artifacts, MA plots, probe distributions, and sample comparison boxplots. We summarized transcript intensities using robust multi-array analysis (RMA) [28]. We determined batch normalization was unnecessary after a guided principal components analysis (gPCA) using sample processing date and array plate ID as potential batch effect confounders [29]. Lastly, we collapsed HTA2.0 transcripts into gene level measurements using the ‘collapseRowsQ’ function with the “maxmean” method from the R package WGCNA [30]. We used the pd.hta.2.0 platform design file (version 3.12.1) and the Bioconductor package “hta20sttranscriptcluster.db” (version 8.3.1) to map manufacturer transcript IDs to genes. We performed all preprocessing steps using R version 3.2.3.

### Western Blotting

Prior to sodium dodecyl sulfate polyacrylamide gel electrophoresis, protein sample concentration was determined using a Pierce BCA Protein Assay Kit (Thermo Scientific). Protein samples were prepared with 1X Laemmli sample buffer (50 mM Tris pH 6.8, 0.02% w/v bromophenol blue, 2% w/v SDS, 10% v/v glycerol, 1% v/v beta-mercaptoethanol, 12.5 mM EDTA) and 50 μg of tumor protein. Volumes were normalized with RIPA buffer including the protease/phosphatase inhibitors described above. SDS-PAGE was performed using a 4-15% Mini-PROTEAN TGX gel (Bio-Rad) for 1 hour at 120V. The samples were then transferred to a nitrocellulose membrane for 2 hours and 45 minutes at 400 mA in cold transfer buffer (384 mM glycine, 50 mM Tris, 20% methanol, 0.005% w/v sodium dodecyl sulfate. Following this, the blots were then blocked in 5% w/v BSA or 5% w/v nonfat dry milk in Tris-buffered saline (137 mM NaCl, 2.7 mM KCl, 19 mM Tris, 0.05% v/v Tween 20, pH 7.4) for 25 minutes. Immunoblotting was performed with the following antibodies and conditions (vendor, species, diluent, dilution, incubation time, incubation temperature): anti-NF1 D7R7D (Cell Signaling, rabbit, 2% BSA, 1:1000, overnight, 4°C), anti-tubulin B-1-2-5 (Santa Cruz, mouse, 2% milk, 1:10000, 1 hour, RT), anti-EGFR D38B1 (Cell Signaling, rabbit, 2% milk, 1:1000-1:2000, 1h, RT), p-ERK½ (p44/42 MAPK) #9101 (Cell Signaling, rabbit, 2% BSA, 1:2000, overnight, 4°C), SUZ12 D39F6 #3737 (Cell Signaling, rabbit, 2% milk, 1:1000, overnight, 4°C). Anti-NF1 D7R7D was a kind gift from Cell Signaling Technologies, Inc.

The binding of the primary antibodies was detected by incubation with secondary antibodies goat anti-rabbit HRP 1:20000 or goat anti-mouse HRP 1:10000 (Jackson Immunoresearch Laboratories Inc.) at room temperature in 2% milk in TBST and detection of HRP activity using Pierce ECL Western Blotting substrate (Thermo Scientific), or in the case of *NF1*, SuperSignal West Femto Maximum Sensitivity Substrate (Thermo Scientific). The chemiluminescent signal was captured with MED-B medical x-ray film (Med X Ray Company Inc.). Between primary antibodies, the membrane was stripped twice for 10 minutes at room temperature using a mild stripping buffer containing 1.5% w/v glycine, 0.1% w/v SDS, 1% v/v Tween 20 at pH 2.2 (Abcam). One sample was eliminated due to low yield, and apparent degradation as determined by western blotting (all proteins examined were undetectable with the exception of tubulin, not shown). Densitometry was performed using Li-COR Image Studio Lite 5.0. Briefly, intensity measurements for *NF1* and tubulin were taken using equally-sized regions for all bands. The background was subtracted using the local median intensity from the left and right borders (size=2) of each measurement region. *NF1* values were divided by tubulin intensity to adjust for protein loading. All measurement ratios were then normalized by dividing values by the “U87+PI” measurement for each blot, respectively.

### Reproducibility of Computational Analyses

We provide software with a permissive open source license to reproduce all computational analyses [31]. Ensuring a stable compute environment, we performed all analyses in a Docker image [32]. This image and source code can be used to freely confirm, modify, and build upon this work.

## RESULTS

### Classifier performance

Using 5-fold cross validation across a parameter sweep, we identified optimal hyperparameters at alpha = 0.15 and L1 mixing = 0.1 (Supplementary Figure S1). To assess model performance, we performed 100 random initializations of five-fold cross-validation. These models had mean test area under the receiver operating characteristic curve (AUROC) of 0.77 (95% Quantiles: 0.53 – 0.95) and a mean train AUROC of 0.997 (95% Quantile: 0.98 – 1.00) (Supplementary Figure S2). We repeated this procedure after TDM transformation (Supplementary Figure S3) and achieved comparable results with alpha = 0.15 and l1 mixing = 0.1 (mean test AUROC = 0.77, 95% Quantiles: 0.51 – 0.96; mean train AUROC = 0.998, 95% Quantiles: 0.99 – 1.00) (Figure 1B). Because the validation set was measured by microarray, we used the classifier trained on TDM transformed data to construct our ensemble classifier. We also determined the Cohen’s D effect size estimate for all training and testing partitions across all 5-fold cross validation iterations of the TDM transformed model (Supplementary Fig S4). The classifier consistently and robustly separated *NF1* wildtype and *NF1* inactivated GBM samples with high effect sizes (Training: mean Cohen’s D = 3.07, 95% CI = 2.24 – 4.16; Testing: mean Cohen’s D = 1.27, 95% CI = 0.19 - 2.67).

### Identification and characterization of NF1 deficient glioblastoma tumor samples

We characterized NF1 protein concentrations as well as other molecules involved in RAS signaling in the 12 GBM samples (Figure 2A). Two samples (CB2, 3HQ) had no apparent NF1 protein. Eight other samples had similar or less NF1 signal than the U87-MG NF1-low control (H5M, LNA, YXL, VVN, R7K, TRM, UNY, W31). Two samples (PBH, RIW) had equal or greater NF1 than the positive control, U87-MG + proteasome inhibitors (preventing NF1 degradation). We also observed variable EGFR content in these samples, with non-existent to low levels (3HQ, YXL, R7K), or medium to large EGFR signal (CB2, H5M, PBH, LNA, YXL, VVN, RIW, TRM, UNY, W31). All GBM samples had high concentrations of phospho-ERK1/2 signal relative to cell line controls. Samples with increased phospho-ERK1/2 may have greater Ras pathway activation. This can be attributed to multiple factors, including increased EGFR expression and/or NF1 inactivation.

Our ensemble classifier predicted four samples to have NF1 inactivation (CB2, UNY, R7K, and 3HQ) and eight samples to be NF1 wildtype (W31, TRM, PBH, VVN, LNA, RIW, H5M, and YXL) (Figure 2B). Because two samples, (CB2 and H5M) were measured on both western blots (Figure 2C), we used the mean of their NF1 protein level across both experiments.

We performed a one-tailed Welch’s t-test to determine if NF1 protein concentrations were significantly higher in *NF1* wildtype versus *NF1* deficient samples based on our classifier predictions (Figure 2D). We did not observe a significant difference across groups (t = −1.38, *p* = 0.098, effect size = 0.699). Additionally, while the effect size was fairly large, a power analysis indicated that 22 samples per group would be required to achieve a power = 0.8 at that effect size. With a lack of glioblastoma samples with quantified NF1 protein available, the trend of less protein present in samples scored as *NF1* inactivated by the classifier nevertheless remains promising.

One of the samples predicted to be NF1 inactive contains detectable NF1 protein (R7K), suggesting that this sample may have NF1 inactivation not detectable by assaying protein, have a different alteration that phenocopies NF1 loss, or is incorrectly predicted by the classifier. Conversely, there are three samples predicted to be NF1 wildtype that have low or undetectable protein (YXL, VVN, W31), which either indicates unknown elements that confound the detection of some NF1 dysregulated tumors or a classification error.

### Highly Contributing Genes

We observed several genes that consistently contributed to the ensemble classifier performance (Figure 3). Since we applied several classifiers to the validation set as an ensemble, we took the sum of all classifier’s gene weights across all 500 iterations to define these consistently contributing genes. While the data indicate that these genes have an impact on classifier performance, the data do not indicate whether changes in the expression of these genes are a direct consequence in changes in NF1 signaling. Expression of genes such as *TXNIP, ARRDC4, ISPD, C10orf107*, and *DUSP18* appear to be predictive of intact NF1 signaling. Among the list of genes that appear to be expressed in tumors with loss of NF1 function are *QPRT, ATF5, HUS1B, PEG10, HMGA2, RSL1D1, and NRG1*. A full list of positive and negative weight genes that were two standard deviations beyond the gene weight distribution is provided in Supplementary Table S2.

**Figure 3.**
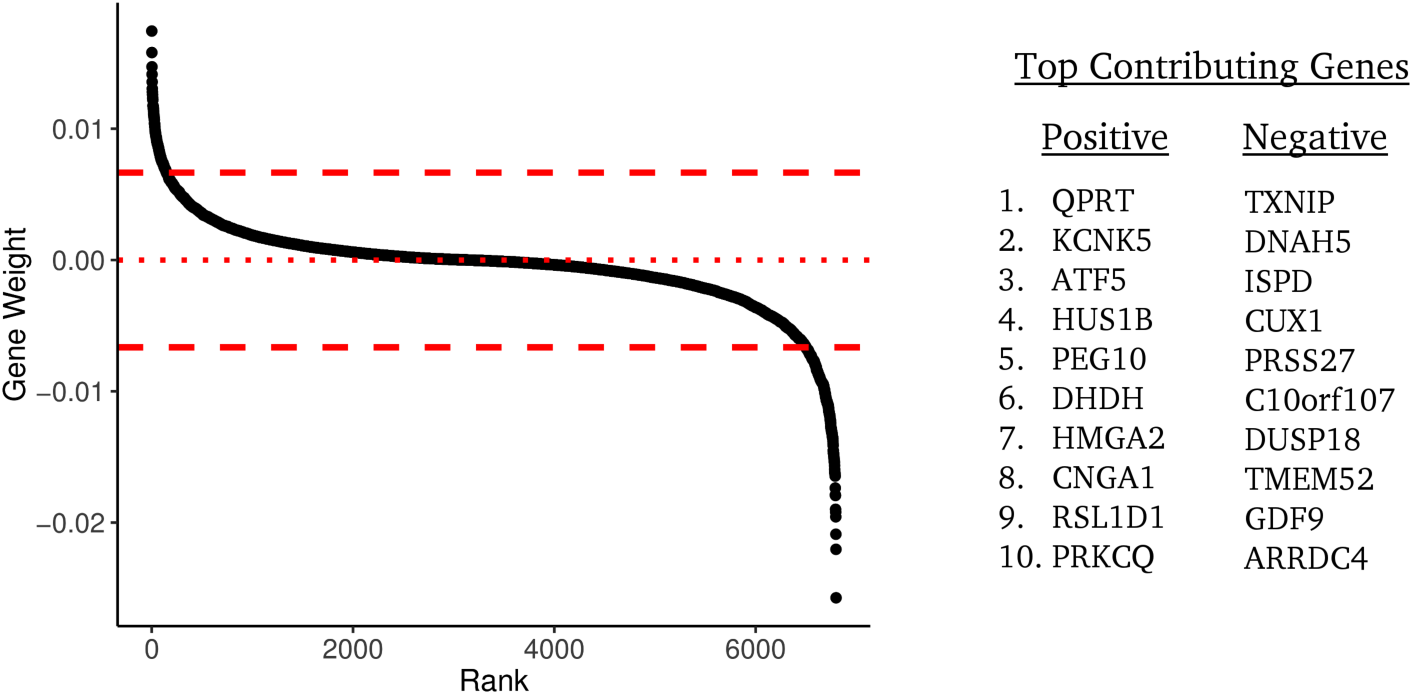
*Genes that contribute to the classifier performance*. Genes are shown ranked by their weighted contribution to the ensemble classifier. Weights are scaled to unit norm. The top 10 positive and top 10 negative contributing high weight genes are given on the right.

We also performed over-representation analysis of the most influential genes in the classifier to identify gene ontology (GO) sets and pathways that may be predictive of NF1 status [33–36]. S3For high-weight genes predictive of intact NF1 signaling, we observed GO sets involved in plasma membrane-localized proteins (GO:0005886, GO:0071944, GO:0016324) and homeostasis (GO:0048871, GO:0001659, GO:0048873, GO:0031224), among others. Annotated pathways associated with genes from this dataset include hematopoietic stem cell differentiation, thyroid cancer, voltage-gated potassium channels, and RHO GTPase functional pathways.

For high-weight genes predictive of NF1 loss of function, we observed GO sets related to cellular adhesion (GO:0007155, GO:0098742), negative regulation of signaling (GO:0009968, GO:0023507, GO:0010648), and nervous system development (GO:0051962, GO0007416, GO: 0050808), among others. These genes were also enriched for elements of the phototransduction cascade and thyroxine production pathways.

## DISCUSSION

A machine learning classifier, based on gene expression data, can capture signal associated with the inactivation of a tumor suppressor. Our classifier is able to detect subtle downstream changes in gene expression as a result of the tumor responding to NF1 loss of function. This finding supports using mRNA as a summary measurement capable of capturing system-wide responses to molecular events beyond transcription factor alterations. Machine learning has been applied to gene expression in a variety of studies with various goals [37–41]. In a similar study, Guinney *et al*. trained a classifier to model RAS activity in colorectal cancer and demonstrated its clinical utility by predicting response to MEK inhibitors and anti-EGFR based treatments [18]. With a wealth of signal embedded in gene expression and a rapidly growing library of datasets, the performance of machine learning models is likely to rapidly improve. An increase in performance leads to more reliable clinical applications that would potentially predict the effectiveness of pathway-specific targeted therapies.

While our classifier was able to predict NF1 inactivation status to an extent, its performance is far from being clinically actionable. A major difficulty in developing a reliable classifier in this case is contamination in gold standard positives and negatives. While we aim to detect NF1 inactivation events, our gold standard positives can only include samples with known *NF1* mutation status. Conversely, we expect that negative samples (about 90% of the data) are also contaminated with NF1 inactivated samples due to protein loss and other mechanisms. We cannot determine scenarios where NF1 is inactivated beyond mutation at scale in the TCGA data. Another challenge with the construction of classifiers from such data is overfitting. Even after hyperparameter optimization we observed substantial overfitting (Figure 2), which has also been observed in competitions (see, for example, supplementary figure S2 of Noren *et al*. 2016 [42] in which the best performing algorithms also overfit). Finally with a small number of positive examples the model performance is unstable, which demonstrates high variability in gold standard samples used to train the model [43]. We employed ensemble classification to mitigate this issue as averaging over heterogeneous models would result in a relatively stable classifier (see Figure 2B). In summary, our results are promising but these challenges are substantial and significant work remains to reach a robust classifier with clinical utility.

The performance of the classifier appears to be impacted by many cancer related genes. For example, genes such as *TXNIP* and *ARRDC4*, which are both indicative of lactic acidosis, correlate with better clinical outcomes, and contribute to predicting tumors with intact NF1 signaling [44]. We also observed transcripts that are more highly expressed in brain tissue than either other normal tissue (*ISPD, C10orf107*), or more highly expressed in normal brain tissue than glioma (*EPHA5*)[45–47]. *DUSP18* contributes to the prediction of *NF1* wildtype status and is a negative regulator of ERK phosphorylation, possibly by regulating *SHP2* phosphorylation [48]. It is unclear whether the expression of these genes is a direct result of *NF1* expression, the result of signaling downstream of NF1, or a consequence of other phenomena (such as expression of *SPRED1*, an NF1 binding partner that is essential for *NF1* signaling). Future studies could elucidate the potential connections between *NF1* and the genes identified as important for the performance of this classifier.

Over-representation analysis of these data highlighted changes in potassium channel expression. It was previously demonstrated that *NF1* wild-type Schwann cells have altered K+ channel activity as compared to *NF1^-/-^* Schwann cells suggesting that this may be one factor by which *NF1* mutant and wild¬type cells can be distinguished [49].

Regarding prediction of NF1 inactivated tumors, we observed several genes that have been linked to cancer such as *QPRT*, which is highly expressed in malignant pheochromocytomas as compared to benign; *RSL1D1* (CSIG), which stabilizes *c-myc* in hepatocellular carcinoma; *PPEF*, which is highly expressed in astrocytic gliomas as compared to normal brain tissue [50–52]; and *PEG10*, a poor prognostic marker and regulator of proliferation, migration, and invasion in several tumor types [53–55]. We also observed *ATF5*, a gene for which expression in malignant glioma is correlated with poor survival [56]. Knockdown of *ATF5* in GBM cells causes cell death *in vitro* and *in vivo* [57]. Analysis of genes that contribute to the prediction of *NF1* inactivation yielded several GO terms related to neural development. It is well established that loss of *NF1* can result in abnormal neural development and/or tumorigenesis [14,58,59]. We also observed genes associated with the mesodermal commitment pathway, components of which are linked to the epithelial to mesenchymal transition in human cancer cells [60–62]. Analysis of this pathway may be informative in identifying tumors with *NF1* loss because mesenchymal GBMs are enriched for tumors with *NF1* loss [63].

Our ensemble classifier was able to robustly detect the samples with the highest and lowest NF1 protein concentrations, but it struggled with samples of intermediate NF1 concentrations. This could be a result of an enrichment of mechanisms causing NF1 inactivation beyond protein abundance, an overrepresentation of mesenchymal tumors in NF1 inactivated samples contaminating dataset splits [63], poor classifier generalizability, or incomplete data transformation between RNAseq and microarray data. Because training and testing performance were similar between transformed and non-transformed data (see Figure 1 and Supplementary Fig S2) we don–t anticipate performance to be impacted much by platform differences or classifier generalizability. Nevertheless, we demonstrated the ability of system-wide gene expression measurements to capture downstream consequences of a complex biological mechanism that would otherwise require several different types of data acquisition to capture.

## CONCLUSIONS

A machine learning classifier for transcriptomic data was able to detect signal associated with the inactivation of NF1, a tumor suppressor gene. The gene is an important regulator of the oncogene *RAS* and is inactivated frequently in GBM and in other tumors. The measurement of NF1 inactivity cannot be comprehensively captured by any single genomic characterization such as targeted sequencing or fluorescence in situ hybridization. This difficulty arises from diverse and complex biological mechanisms that inactivate the tumor suppressor in a variety of ways. However, we demonstrated that measuring system-wide RNA can capture subtle downstream changes that occur in response to NF1 inactivation. Improving classification performance is required before transitioning such a model into clinical use, but our method could be used to characterize cell lines or patient-derived xenograft (PDX) models with inactive NF1. Eventually, with more data and improved classification, we expect machine-learning models constructed on system-wide transcriptomics will translate into clinically relevant predictions that will guide targeted therapy.

## DECLARATIONS

### Ethics approval and consent to participate

The Dartmouth-Hitchcock Medical Center Committee for the Protection of Human Subjects declared IRB review and approval is not required since the study does not meet the regulatory definition of human subjects.

### Consent for publication

Not Applicable

### Availability of data and materials

All source code is available under a permissive open source license in the nf1_inactivation GitHub repository, https://github.com/greenelab/nf1 inactivation. We also provide a docker image to replicate the computational environment at https://hub.docker.com/r/gregway/nf1_inactivation/. Additionally, all validation data is available under Gene Expression Omnibus accession number GSE85033.

## Author Contributions

GPW built, analyzed, and interpreted the classifier, generated the figures, created all source code, and wrote the manuscript. RJA performed all experiments, interpreted the classifier, and wrote the manuscript. SJB interpreted the results and was a major contributor to the manuscript. CEF was a major contributor to the study design and was a major contributor to the manuscript. YS designed the study, interpreted the results, and wrote the manuscript. CSG designed the study, interpreted the results, and wrote the manuscript. All authors read and approved the final manuscript.

## Acknowledgements

This work was supported by the MMC BioBank, a core facility of the Maine Medical Center Research Institute, and the Dartmouth Genomics Shared Resource, a core facility of the Norris Cotton Cancer Center.

## SUPPLEMENTARY FIGURES

**Supplementary Figure S1.**
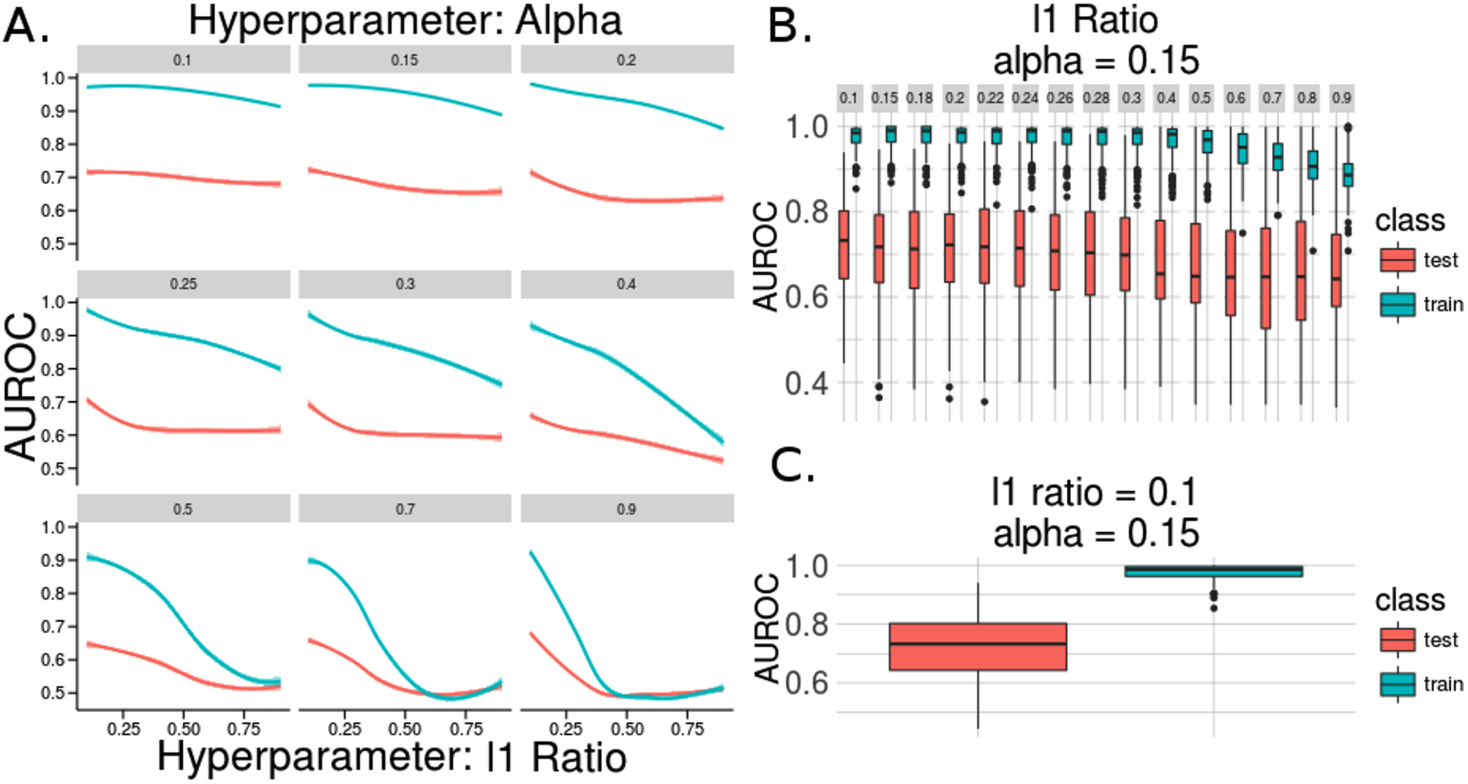
Non-transformed RNAseq results of The Cancer Genome Atlas Glioblastoma parameter sweep for stochastic gradient descent logistic classifiers with elastic net penalty. (A) Training and testing area under the receiver operating characteristic curve (AUROC) ass is given for each parameter tested. All accuracies are presented following 5-fold cross validation after 50 random initializations. (B) The l1 mixing parameter with the optimal alpha and (C) the classifier performance across all random starts for the best hyperparameters.

**Supplementary Figure S2.**
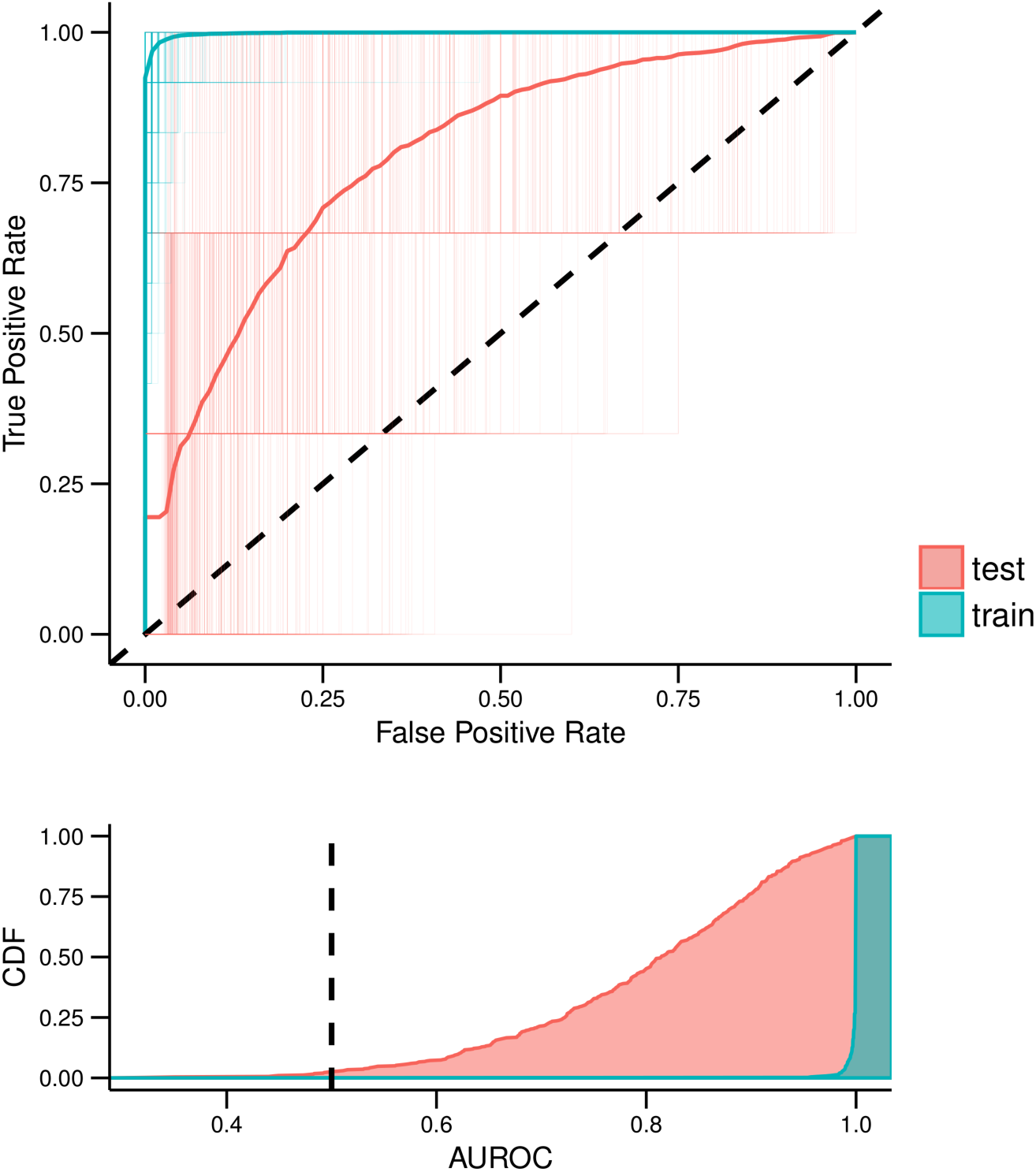
Logistic regression classifier with elastic net penalty training and testing errors over 100 iterations for non-transformed The Cancer Genome Atlas Glioblastoma RNAseq data. (A) Receiver operating characteristic (ROC) curve and shows the average training and testing performance of 5-fold cross validation over 100 random initializations as well as each individual classifier in the ensemble model. (B) The cumulative density of area under the ROC curve (AUROC) for all training and testing partitions.

**Supplementary Figure S3.**
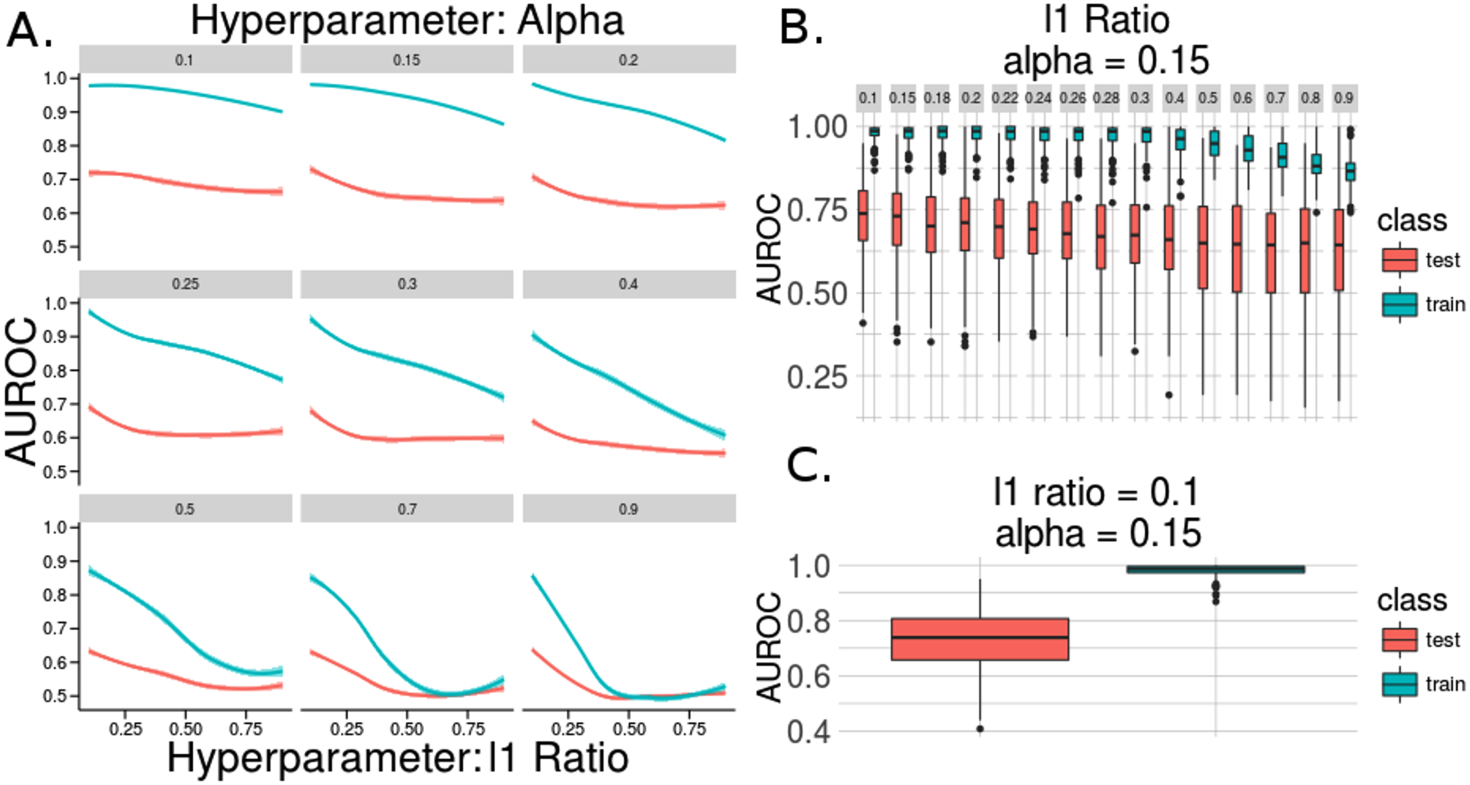
Training Distribution Matching (TDM) transformation of RNAseq results of The Cancer Genome Atlas Glioblastoma parameter sweep for stochastic gradient descent logistic classifier with elastic net penalty. (A) Training and testing area under the receiver operating characteristic curve (AUROC) is given for each parameter tested. All accuracies are presented following 5-fold cross validation after 100 random initializations. (B) The l1 mixing parameter with the optimal alpha and (C) the classifier performance across all random starts for the best hyperparameters.

**Supplementary Figure S4.**
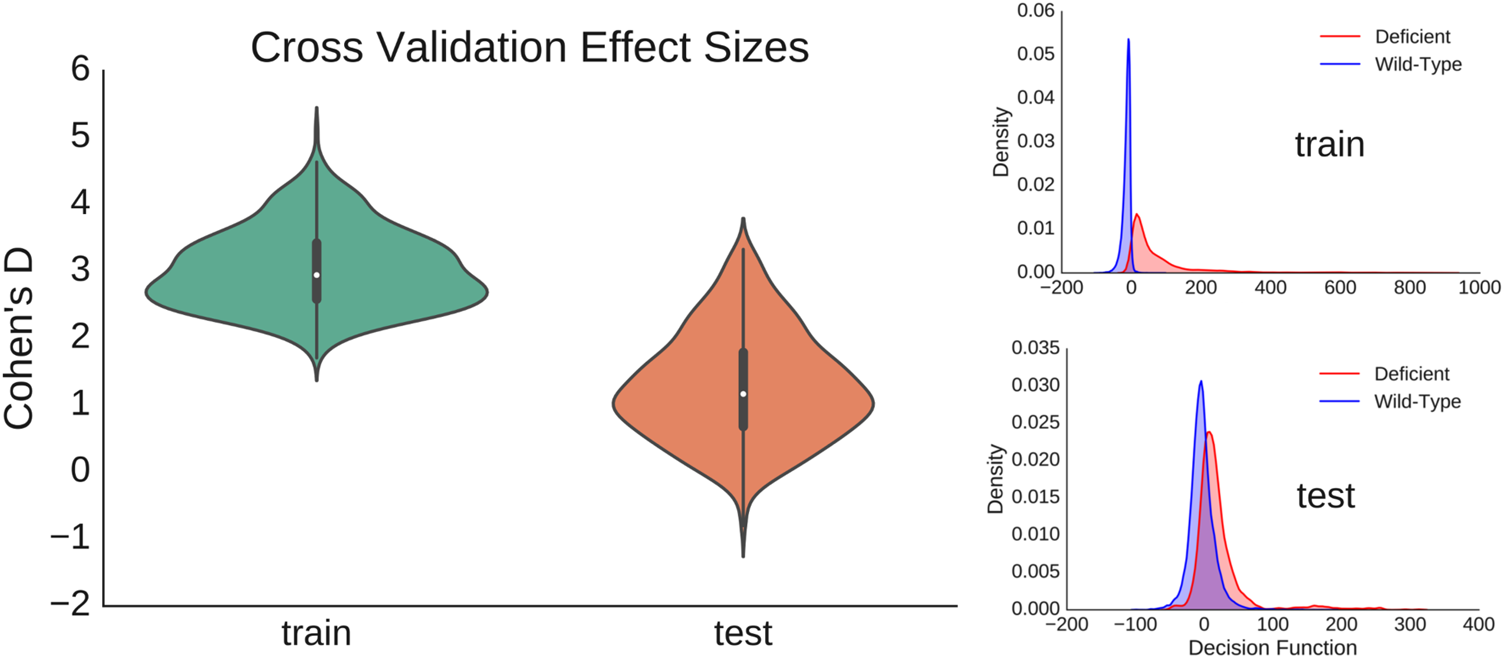
Cohen’s D effect size estimates across 5 fold cross validation parameters for all 100 iterations of the TDM transformed ensemble classifier. The effect size for the test set is consistently lower than the training set (left). Additionally, the training and testing decision functions for gold standard NF1 deficient vs. NF1 wildtype samples shows a difference in mean estimates (right). The decision function represents the raw score of all samples as applied to the respective classifiers through each of the 100 iterations of 5 fold cross validation on the TCGA training set.

## References

1. Martin GA, Viskoohil D, Bollag G, McCabe PC, Crosier WJ, Haubruck H, et al. The GAP-related domain of the neurofibromatosis type 1 gene product interacts with ras p21. Cell. 1990;63:843–849.

2. Xu G, O’Connell P, Viskochil D, Cawthon R, Robertson M, Culver M, et al. The neurofibromatosis type 1 gene encodes a protein related to GAP. Cell. 1990;62:599–608.

3. Boyd KP, Korf BR, Theos A. Neurofibromatosis type 1. J. Am. Acad. Dermatol. 2009;61:1–14.

4. Dogra B, Rana K. Facial plexiform neurofibromatosis: A surgical challenge. Indian Dermatol. Online J. 2013;4:195.

5. Evans DGR, Baser M, McGaughran J, Sharif S, Howard E, Moran A. Malignant peripheral nerve sheath tumours in neurofibromatosis 1. J. Med. Genet. 2002;39:311–4.

6. Rad E, Tee AR. Neurofibromatosis type 1: Fundamental insights into cell signalling and cancer. Semin. Cell Dev. Biol. 2016;52:39–46.

7. Ratner N, Miller SJ. A RASopathy gene commonly mutated in cancer: the neurofibromatosis type 1 tumour suppressor. Nat. Rev. Cancer. 2015;15:290–301.

8. Wood M, Rawe M, Johansson G, Pang S, Soderquist RS, Patel AV, et al. Discovery of a Small Molecule Targeting IRA2 Deletion in Budding Yeast and Neurofibromin Loss in Malignant Peripheral Nerve Sheath Tumor Cells. Mol. Cancer Ther. 2011;10:1740–50.

9. Allaway RJ, Fischer DA, de Abreu FB, Gardner TB, Gordon SR, Barth RJ, et al. Genomic characterization of patient-derived xenograft models established from fine needle aspirate biopsies of a primary pancreatic ductal adenocarcinoma and from patient-matched metastatic sites. Oncotarget. 2016;7:17087–102.

10. McGillicuddy LT, Fromm JA, Hollstein PE, Kubek S, Beroukhim R, De Raedt T, et al. Proteasomal and Genetic Inactivation of the NF1 Tumor Suppressor in Gliomagenesis. Cancer Cell. 2009;16:44–54.

11. Subramanian S, Thayanithy V, West RB, Lee C-H, Beck AH, Zhu S, et al. Genome-wide transcriptome analyses reveal p53 inactivation mediated loss of miR-34a expression in malignant peripheral nerve sheath tumours. J. Pathol. 2010;220:58–70.

12. Wallace MR, Andersen LB, Saulino AM, Gregory PE, Glover TW, Collins FS. A de novo Alu insertion results in neurofibromatosis type 1. Nature 1991;353:864–6.

13. Skuse GR, Cappione AJ, Sowden M, Metheny LJ, Smith HC. The Neurofibromatosis Type I Messenger RNA Undergoes Base-Modification RNA Editing. Nucleic Acids Res. 1996;24:478–86.

14. Cichowski K, Jacks T. NF1 tumor suppressor gene function: narrowing the GAP. Cell. 2001;104:593–604.

15. UCSC Xena [Internet]. Available from: http://xena.ucsc.edu/

16. Brennan CW, Verhaak RGW, McKenna A, Campos B, Noushmehr H, Salama SR, et al. The Somatic Genomic Landscape of Glioblastoma. Cell. 2013;155:462–77.

17. Thompson JA, Tan J, Greene CS. Cross-platform normalization of microarray and RNA-seq data for machine learning applications. PeerJ. 2016;4:e1621.

18. Guinney J, Ferte C, Dry J, McEwen R, Manceau G, Kao K, et al. Modeling RAS Phenotype in Colorectal Cancer Uncovers Novel Molecular Traits of RAS Dependency and Improves Prediction of Response to Targeted Agents in Patients. Clin. Cancer Res. 2014;20:265–72.

19. Liang Y, Liu C, Luan X-Z, Leung K-S, Chan T-M, Xu Z-B, et al. Sparse logistic regression with a L1/2 penalty for gene selection in cancer classification. BMC Bioinformatics. 2013;14:198.

20. Sokolov A, Carlin DE, Paull EO, Baertsch R, Stuart JM. Pathway-Based Genomics Prediction using Generalized Elastic Net. Przytycka TM, editor. PLOS Comput. Biol. 2016;12:e1004790.

21. Pedregosa F, Varoquaux G, Gramfort A, Michel V, Thirion B, Grisel O, et al. Scikit-learn: Machine Learning in Python. CoRR. 2012;

22. Cohen J. Statistical power analysis for the behavioral sciences. New York: Academic Press; 1969.

23. Allen M, Bjerke M, Edlund H, Nelander S, Westermark B. Origin of the U87MG glioma cell line: Good news and bad news. Sci. Transl. Med. 2016;8:354re3–354re3.

24. Daginakatte GC, Gutmann DH. Neurofibromatosis-1 (Nf1) heterozygous brain microglia elaborate paracrine factors that promote Nf1-deficient astrocyte and glioma growth. Hum. Mol. Genet. 2007;16:1098–112.

25. Hollstein PE, Cichowski K. Identifying the Ubiquitin Ligase Complex That Regulates the NF1 Tumor Suppressor and Ras. Cancer Discov. 2013;3:880–93.

26. Tan X, Wang S, Yang B, Zhu L, Yin B, Chao T, et al. The CREB-miR-9 Negative Feedback Minicircuitry Coordinates the Migration and Proliferation of Glioma Cells. Li J, editor. PLoS ONE. 2012;7:e49570.

27. Carvalho BS, Irizarry RA. A framework for oligonucleotide microarray preprocessing. Bioinforma. Oxf. Engl. 2010;26:2363–7.

28. Irizarry RA, Bolstad BM, Collin F, Cope LM, Hobbs B, Speed TP, Summaries of Affymetrix GeneChip probe level data. Nucleic Acids Res. 2003;31:e15.

29. Reese SE, Archer KJ, Therneau TM, Atkinson EJ, Vachon CM, de Andrade M, et al. A new statistic for identifying batch effects in high-throughput genomic data that uses guided principal component analysis. Bioinformatics. 2013;29:2877–83.

30. Miller JA, Cai C, Langfelder P, Geschwind DH, Kurian SM, Salomon DR, et al. Strategies for aggregating gene expression data: The collapseRows R function. BMC Bioinformatics. 2011;12:322.

31. Greg Way. nf1_inactivation: Pre-Release. 2016 [cited 2016 Aug 1]; Available from: http://dx.doi.org/10.5281/zenodo.58864

32. Boettiger C. An introduction to Docker for reproducible research. ACM SIGOPS Oper. Syst. Rev. 2015;49:71–9.

33. Kamburov A, Wierling C, Lehrach H, Herwig R. ConsensusPathDB––a database for integrating human functional interaction networks. Nucleic Acids Res. 2009;37:D623–8.

34. Kamburov A, Stelzl U, Lehrach H, Herwig R. The ConsensusPathDB interaction database: 2013 update. Nucleic Acids Res. 2013;41:D793–800.

35. Ashburner M, Ball CA, Blake JA, Botstein D, Butler H, Cherry JM, et al. Gene Ontology: tool for the unification of biology. Nat. Genet. 2000;25:25–9.

36. The Gene Ontology Consortium. Gene Ontology Consortium: going forward. Nucleic Acids Res. 2015;43:D1049–56.

37. Molla M, Waddell M, Page D, Shavlik J. Using Machine Learning to Design and Interpret Gene-Expression Microarrays. AI Mag. 2004;25:23–44.

38. Bastani M, Vos L, Asgarian N, Deschenes J, Graham K, Mackey J, et al. A Machine Learned Classifier That Uses Gene Expression Data to Accurately Predict Estrogen Receptor Status. Rogers S, editor. PLoS ONE. 2013;8:e82144.

39. Pirooznia M, Yang JY, Yang MQ, Deng Y. A comparative study of different machine learning methods on microarray gene expression data. BMC Genomics. 2008;9:S13.

40. Chou W-C, Ma Q, Yang S, Cao S, Klingeman DM, Brown SD, et al. Analysis of strand-specific RNA-seq data using machine learning reveals the structures of transcription units in Clostridium thermocellum. Nucleic Acids Res. 2015;43:e67–e67.

41. Tan J, Hammond JH, Hogan DA, Greene CS. ADAGE-Based Integration of Publicly Available *Pseudomonas aeruginosa* Gene Expression Data with Denoising Autoencoders Illuminates Microbe-Host Interactions. Gilbert JA, editor. mSystems. 2016;1:e00025–15.

42. Noren DP, Long BL, Norel R, Rrhissorrakrai K, Hess K, Hu CW, et al. A Crowdsourcing Approach to Developing and Assessing Prediction Algorithms for AML Prognosis. Tan K, editor. PLOS Comput. Biol. 2016;12:e1004890.

43. Yu B. Stability. Bernoulli. 2013;19:1484–500.

44. Chen JL-Y, Merl D, Peterson CW, Wu J, Liu PY, Yin H, et al. Lactic Acidosis Triggers Starvation Response with Paradoxical Induction of TXNIP through MondoA. Gibson G, editor. PLoS Genet. 2010;6:e1001093.

45. Willer T, Lee H, Lommel M, Yoshida-Moriguchi T, de Bernabe DBV, Venzke D, et al. ISPD loss-of-function mutations disrupt dystroglycan O-mannosylation and cause Walker-Warburg syndrome. Nat. Genet. 2012;44:575–80.

46. Thierry-Mieg D, Thierry-Mieg J. AceView: a comprehensive cDNA-supported gene and transcripts annotation. Genome Biol. 2006;7 Suppl 1:S12.1–14.

47. Almog N, Ma L, Raychowdhury R, Schwager C, Erber R, Short S, et al. Transcriptional Switch of Dormant Tumors to Fast-Growing Angiogenic Phenotype. Cancer Res. 2009;69:836–44.

48. Sacco F, Boldt K, Calderone A, Panni S, Paoluzi S, Castagnoli L, et al. Combining affinity proteomics and network context to identify new phosphatase substrates and adapters in growth pathways. Front. Genet. [Internet]. 2014 [cited 2016 Aug 1];5. Available from: http://journal.frontiersin.org/article/10.3389/fgene.2014.00115/abstract

49. Xu Y, Chiamvimonvat N, Vázquez AE, Akunuru S, Ratner N, Yamoah EN. Gene-targeted deletion of neurofibromin enhances the expression of a transient outward K+ current in Schwann cells: a protein kinase A-mediated mechanism. J. Neurosci. Off. J. Soc. Neurosci. 2002;22:9194–202.

50. Thouënnon E, Elkahloun AG, Guillemot J, Gimenez-Roqueplo A-P, Bertherat J, Pierre A, et al. Identification of Potential Gene Markers and Insights into the Pathophysiology of Pheochromocytoma Malignancy. J. Clin. Endocrinol. Metab. 2007;92:4865–72.

51. Cheng Q, Yuan F, Lu F, Zhang B, Chen T, Chen X, et al. CSIG promotes hepatocellular carcinoma proliferation by activating c-MYC expression. Oncotarget. 2015;6:4733–44.

52. Bageritz J, Puccio L, Piro RM, Hovestadt V, Phillips E, Pankert T, et al. Stem cell characteristics in glioblastoma are maintained by the ecto-nucleotidase E-NPP1. Cell Death Differ. 2014;21:929–40.

53. Deng X, Hu Y, Ding Q, Han R, Guo Q, Qin J, et al. PEG10 plays a crucial role in human lung cancer proliferation, progression, prognosis and metastasis. Oncol. Rep. 2014;32:2159–67.

54. Li C-M, Margolin AA, Salas M, Memeo L, Mansukhani M, Hibshoosh H, et al. PEG10 is a c-MYC target gene in cancer cells. Cancer Res. 2006;66:665–72.

55. Akamatsu S, Wyatt AW, Lin D, Lysakowski S, Zhang F, Kim S, et al. The Placental Gene PEG10 Promotes Progression of Neuroendocrine Prostate Cancer. Cell Rep. 2015;12:922–36.

56. Sheng Z, Li L, Zhu LJ, Smith TW, Demers A, Ross AH, et al. A genome-wide RNA interference screen reveals an essential CREB3L2-ATF5-MCL1 survival pathway in malignant glioma with therapeutic implications. Nat. Med. 2010;16:671–7.

57. Greene LA, Lee HY, Angelastro JM. The transcription factor ATF5: role in neurodevelopment and neural tumors. J. Neurochem. 2009;108:11–22.

58. Zhu Y, Romero MI, Ghosh P, Ye Z, Charnay P, Rushing EJ, et al. Ablation of NF1 function in neurons induces abnormal development of cerebral cortex and reactive gliosis in the brain. Genes Dev. 2001;15:859–76.

59. Joseph NM, Mosher JT, Buchstaller J, Snider P, McKeever PE, Lim M, et al. The Loss of Nf1 Transiently Promotes Self-Renewal but Not Tumorigenesis by Neural Crest Stem Cells. Cancer Cell. 2008;13:129–40.

60. Morishita A, Zaidi MR, Mitoro A, Sankarasharma D, Szabolcs M, Okada Y, et al. HMGA2 is a driver of tumor metastasis. Cancer Res. 2013;73:4289–99.

61. de Boeck M, Cui C, Mulder AA, Jost CR, Ikeno S, ten Dijke P. Smad6 determines BMP-regulated invasive behaviour of breast cancer cells in a zebrafish xenograft model. Sci. Rep. 2016;6:24968.

62. Salomonis N, Mshel016, Cirillo E, Hanspers K, Kutmon M. Mesodermal Commitment Pathway (Homo sapiens). http://www.wikipathways.org/index.php/Pathway:WP2857.

63. Verhaak RGW, Hoadley KA, Purdom E, Wang V, Qi Y, Wilkerson MD, et al. Integrated genomic analysis identifies clinically relevant subtypes of glioblastoma characterized by abnormalities in PDGFRA, IDH1, EGFR, and NF1. Cancer Cell. 2010;17:98–110.

